# Early-life adversity alters the adult blood proteome primarily in female mice

**DOI:** 10.64898/2026.09.28.755115

**Authors:** Madison M. Garvin, Ae Ra Kim, Jessica L. Bolton, Yun Liang

## Abstract

**Background:** Early-life adversity (ELA) is one of the strongest predictors of poor mental and physical health later in life, increasing an individual’s risk for disorders ranging from depression to metabolic syndrome. However, whether and how ELA shapes immune memory through adulthood, subsequently affecting susceptibility to immune-associated diseases such as autoimmunity, remains elusive.

**Methods:** To address this question, we employed a translationally and ethologically relevant mouse model of resource deprivation, the limited bedding and nesting model, to study the long-term effects of ELA on the immune system. ELA was induced by limited bedding and nesting material in the cage during postnatal days (P)2-9, which provokes fragmented and unpredictable maternal behavior in the dams, and results in chronic stress in the pups. Nucleic acid Linked ImmunoSandwich Assay (NULISA) was used to survey the proteome of adult (∼P90) mice of both sexes that experienced ELA vs. control conditions.

**Results:** Using a novel proteomic method, we detected significant upregulation in the levels of IL-17A in the adult blood, primarily in females. Within females, ELA also upregulated IL-13 and CCL3. Three proteins, HGF, PDGFA, and WNT16, were significantly downregulated in ELA mice of both sexes.

**Conclusions:** Being a hallmark proinflammatory cytokine, IL-17A is found to increase in patients with systemic lupus erythematosus (SLE), as well as in animals with lupus-like diseases in virtually all reports. Importantly, autoimmune diseases like SLE have a higher prevalence in women than men, and our data support this sex bias. Together, our data suggest that ELA shapes long-term immune function in a way that may increase risk for autoimmune diseases, with promise for the development of more targeted and personalized therapies for autoimmunity.

## 1. Introduction

Early-life adversity (ELA) is one of the strongest predictors of poor mental and physical health later in life, increasing an individual’s risk for disorders ranging from depression to metabolic syndrome. ELA involves severe deprivation and/or situations that will require significant physiological or psychological adaptations during childhood or adolescence^1^, when there is increased plasticity during sensitive periods of development. One putative mechanism of ELA’s effects on later health is inflammation, which has been previously explored primarily in the context of depression. ELA is associated with elevated markers of inflammation such as CRP, IL-6, and TNF-alpha^2^, and anti-inflammatory drugs have been shown to decrease depressive symptoms in depressed patients with increased baseline levels of inflammation^3^. Furthermore, there is now ample evidence for brain-immune crosstalk in the normal developing brain, which can be perturbed by ELA to result in “rewiring” of brain circuits that results in a long-term increase in depression risk. For example, ELA inhibits the function of microglia, the resident macrophages of the brain, during the critical period for synaptic pruning, resulting in an excess of excitatory synapses, and later hyperexcitability of stress-responsive brain regions, and ultimately aberrant stress responses and increased risk for depression^4^. Despite these lines of evidence suggesting brain-immune crosstalk, whether ELA is a direct trigger for immune dysregulation in the periphery and its molecular mechanism remains elusive.

Autoimmune diseases are a group of chronic disorders that involves one’s immune system attacking self-organs^2^. As a category, autoimmune diseases affect 50 million Americans, are the second highest cause of chronic illness, and the top causes of morbidity in women in the U.S^2^. The exact causes of autoimmune diseases are not known, even for the most-studied disorders such as systemic lupus erythematosus (SLE)^5^. While genome-wide association studies have identified lupus-susceptible genes in NFκB, IFN and additional immune-associated pathways, other studies indicate that genetics only contribute to 30% of autoimmune pathogenesis^6,7^, suggesting that environmental factors are important contributors. Indeed, a recent questionnaire-based study found that childhood trauma increases the risk of developing lupus later in life in women by 2.57 times^8^. Depression is a highly prevalent and significant comorbidity in SLE that also affects more women than men^9^, suggesting a possible convergence in mechanisms programming the brain and body after ELA.

Here, we aimed to test the hypothesis that ELA alters inflammatory markers in adulthood that may increase the risk for autoimmune diseases like lupus. Using a novel proteomic method, we detected significant upregulation in the levels of IL-17A, a hallmark proinflammatory cytokine that is found to increase in patients with SLE and animal models^10–12^, and additional immune-associated proteins in the adult blood, particularly in females. Thus, these data support our hypothesis and establish ELA as a key component of the ‘exposome’ that impacts immune health later in life.

## 2. Materials and Methods

### 2.1 Animals

All mice were housed in quiet, uncrowded conditions on a 12-hr light/dark cycle with *ad libitum* access to food and water. The mice employed in this study were on a C57BL/6 background and included CX3CR1-BAC-Cre^-^::MerTK^fl/fl^ (functionally wild-type mice; CX3CR1-BAC-Cre mice were from GENSAT; MMRRC stock #036395-UCD; obtained from Dr. Staci Bilbo, Duke University; Mertk^fl/fl^ mice were obtained from Dr. Carla Rothlin, Yale University). Both male and female offspring were used throughout the entire study. All experiments were performed in accordance with National Institutes of Health (NIH) guidelines and were approved by the Georgia State University Animal Care and Use Committee.

### 2.2 Limited bedding and nesting (LBN) model of early-life adversity (ELA)

Experimental dams were checked daily for pregnancy and then single-housed in our quiet, temperature-controlled maternal care room. On the morning of postnatal day (P)2, litters were culled to 8 pups and dams and their litters were randomly assigned to either a control or early-life adversity (ELA) group. Dams in the control group were transferred to a standard cage with ∼220 grams of corn cob bedding and one full cotton-square nestlet. Dams in the ELA group were transferred to a cage fitted with a mesh platform (25 x 19 cm, cat # 4700313244; McNichols Co., Atlanta, GA) sitting approximately 2.5 cm above the floor. Bedding was reduced to ∼110 grams and half of a cotton square nestlet was provided^13^. The cages for both groups remained undisturbed from P2 until P10 when the dams and pups were transferred back to standard cages in the colony room for growing up to adulthood for behavioral testing or P10 pups were euthanized to assess excitatory synapses.

### 2.3 Method Details

#### 2.3.1 Blood collection

Adult mice (∼P90) of both sexes were euthanized with Euthasol (390 mg/kg) and perfused transcardially with 1x phosphate-buffered saline (PBS; pH = 7.4) followed by 4% paraformaldehyde in 0.1 sodium phosphate buffer (pH = 7.4). Blood samples (∼100 µl) were collected from the heart prior to insertion of the perfusion needle. period. Following a 30-min period at room temperature to allow for blood clotting, samples were centrifuged at 3000 x *g* for 15 min, and serum was collected and then stored at -20°C until later analysis.

#### 2.3.2 Nucleic acid Linked ImmunoSandwich Assay (NULISA)

Mouse serum was analyzed for multi-protein quantification using the NULISASeq platform following established procedures at Alamar Biosciences^14^. Samples were briefly centrifuged at 10,000× g for 10 min, followed by incubation with the capture and detection antibody cocktails. The formed immunocomplex was captured by the dT beads, washed, and further captured by the streptavidin beads. After additional washing, the reporter molecules were eluted and collected for quantification by next-generation-sequencing. For next generation sequencing analysis, the library was cleaned using the AMpure XP reagent (Monroviao, CA, USA) and quantified before being sequenced using the Illumina NextSeq instrument (San Diego, CA, USA). Each sample was identified using a unique sample barcode in the ligator sequence, normalized and analyzed for protein quantification following established pipelines^14^ .

### 2.4 Quantification and Statistical Analysis

Statistical analyses were all performed using GraphPad Prism 10.0 software (GraphPad, San Diego, CA, USA). Two-way ANOVAs (Sex x ELA) or unpaired *t-*tests (CTL vs. ELA) were used to analyze all data. Any significant interactions of ELA x Sex were followed by Šídák’s post hoc tests. For the sake of consistency, all data were separated by sex in the graphs due to the observation of multiple sex differences. Data are presented as mean ± SEM and Grubbs’ test was used to remove statistical outliers (alpha=0.05) from the data.

All outliers were removed prior to final statistical analysis, and all experimental analyses were conducted in a blinded manner, with the experimenter having no knowledge of experimental group.

## 3. Results

### 3.1 ELA provokes immune changes in the blood of adult mice

To understand whether ELA impacts the peripheral immune system and risk for autoimmune diseases, we obtained sera from 3-month-old mice reared in control or ELA conditions. On the molecular level, we quantified cytokines, chemokines and other secreted proteins in the serum of CTL versus ELA mice. NULISA analysis suggested a separation of female versus male mice in their response to ELA (Figure 1). Intriguingly, in the developing brain, males are more vulnerable to the inhibition of microglial function by ELA^4^, but it appears that the opposite pattern predominates in the periphery in adulthood, where females are more sensitive to the effects of ELA. Three proteins, including HGF, PDGFA and WNT16, were significantly downregulated in the blood of ELA mice compared to control (P < 0.05, Figure 3A). In contrast, IL-17A was identified as the single protein significantly upregulated post-ELA (P < 0.05, Figure 2B). This upregulation of IL-17A was driven by its 3.47-fold increase in females (Figure 2B), further highlighting sex differences in the immune response to ELA.

**Figure 1.**
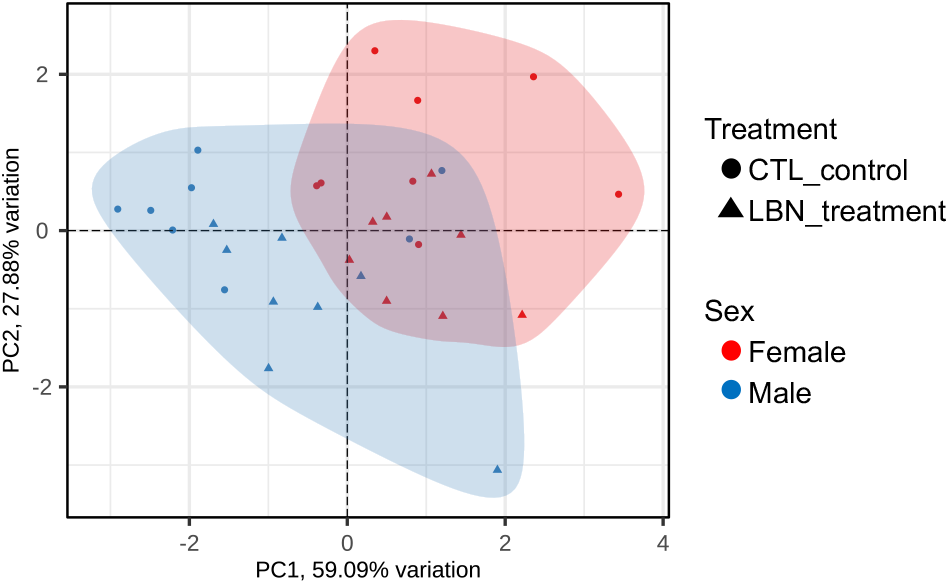
Principal component analysis (PCA) plot of proteome profiles of ELA (LBN-treated) versus control mice. Circles= CTL, control. Triangles= LBN, limited bedding and nesting. Red= females. Blue= males.

**Figure 2.**
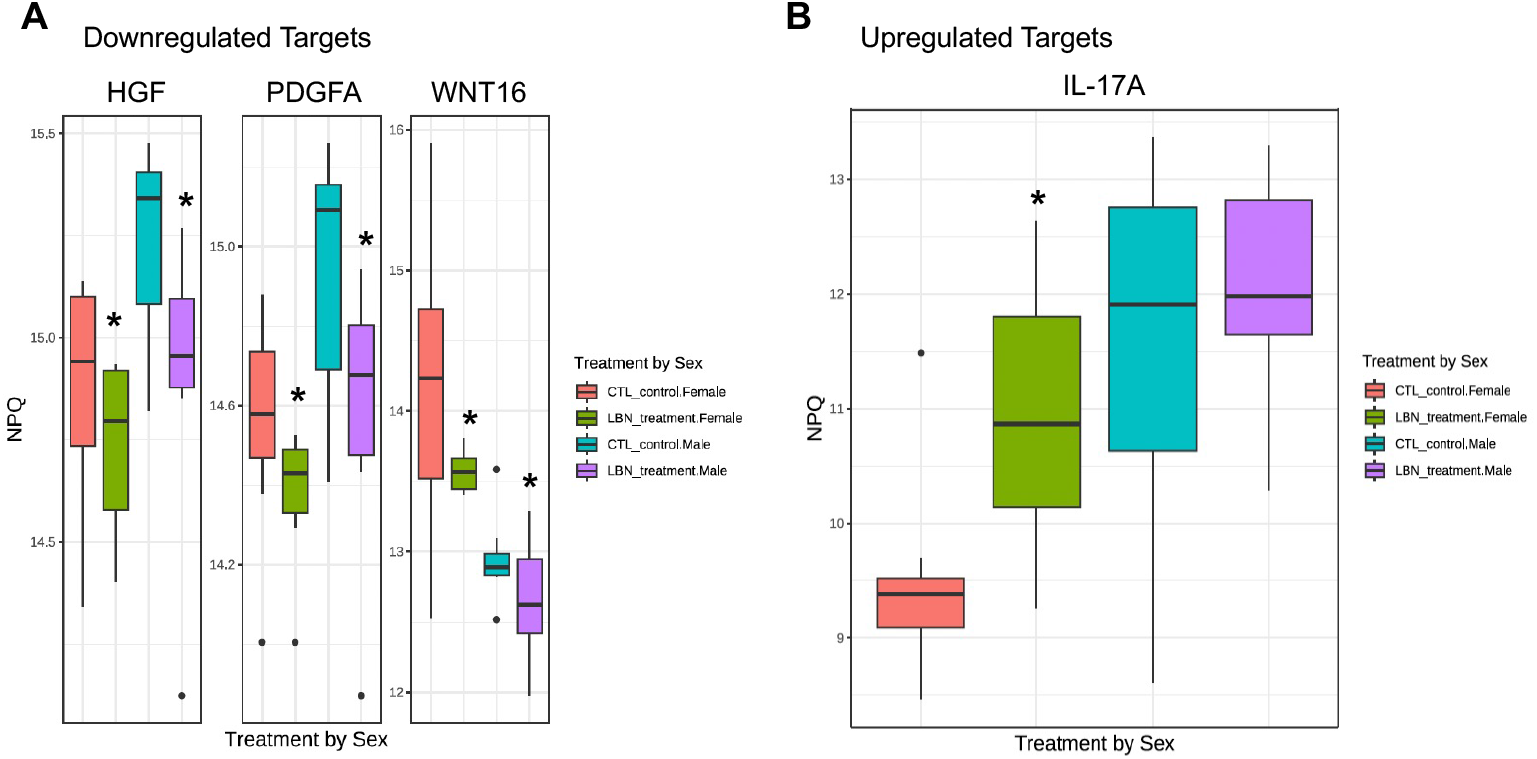
Blood proteins significantly regulated by ELA. Normalized protein quantification (NPQ) levels of ELA-downregulated (A) and upregulated (B) proteins are shown, which indicate expression levels on the log2-scale, are shown for the indicated groups. CTL, control. LBN, limited bedding and nesting (ELA). From left to right, CTL female, LBN female, CTL male, LBN male. *, *p*<0.05, Tukey’s post hoc test indicating significant difference between CTL and LBN for each sex in A, and in females only in B.

**Figure 3.**
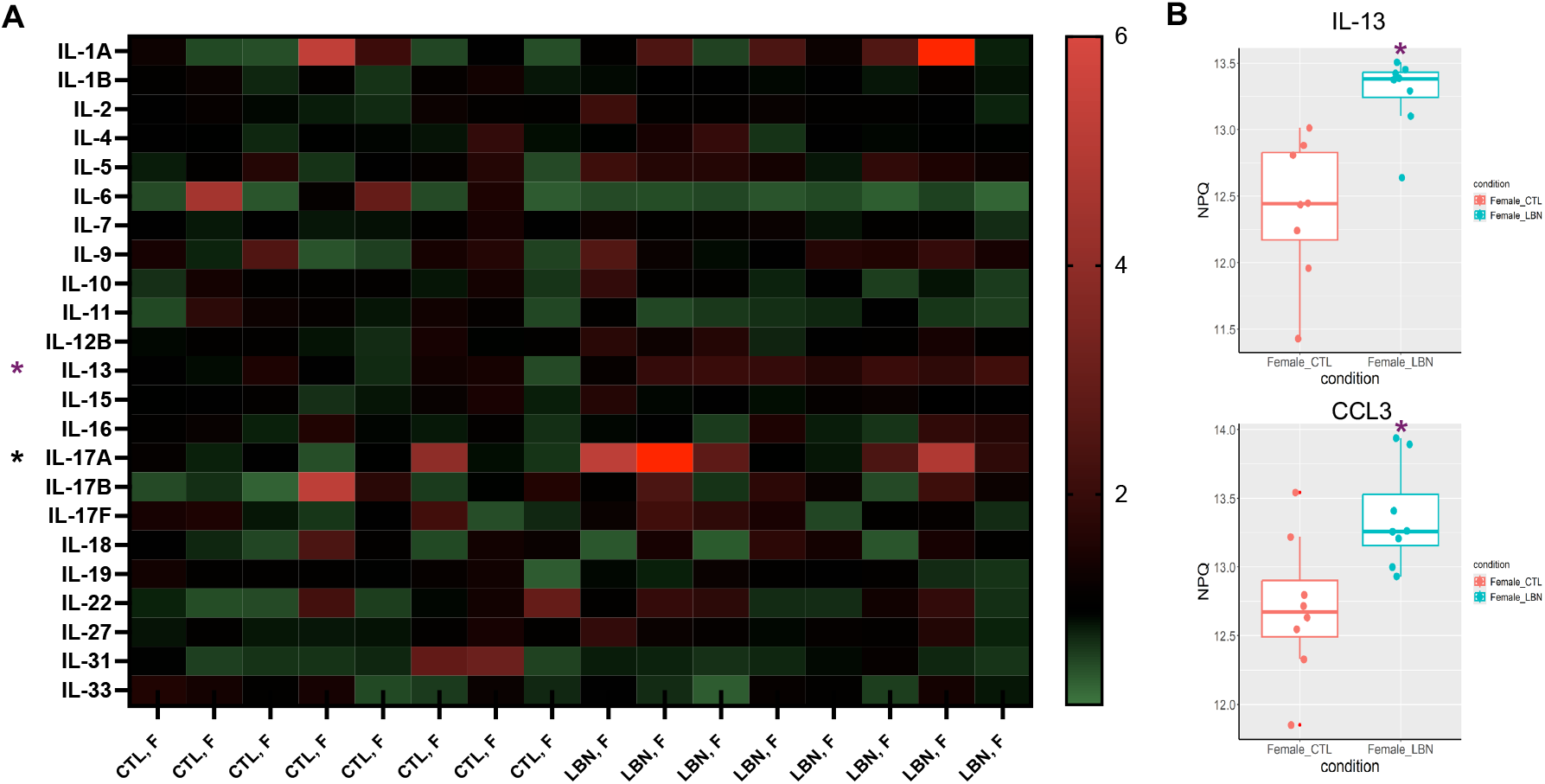
ELA regulation of cytokines in female mice. (a) Heatmap of individual ILs in indicated groups, with scale bar showing the fold-change in protein levels. (b) Normalized protein quantification (NPQ) levels of IL-13 (top) and CCL3 (bottom). CTL, control. LBN, limited bedding and nesting. F, female. *, *p*<0.05, Student’s t-test. * in black, significantly upregulated in females and when both sexes are combined. * in purple, significantly upregulated in females only.

Among the interleukin (IL) family, IL-17A was the most significantly upregulated by ELA in our preliminary data, even when both sexes are combined for the analysis with sex differences removed (Figure 2B). Analyzing each sex separately identified additional ELA-regulated targets such as IL-13 (1.87-fold increase post-ELA in females, Figure 3B) and CCL3 (2.07-fold increase post-ELA in females, Figure 3B).

## 4. Discussion

In a translationally and ethologically relevant rodent model for ELA now employed by hundreds of labs around the world, we have previously demonstrated that exposure to an impoverished environment and unpredictable/fragmented maternal care early in life (due to limited bedding and nesting material) provokes an exaggerated stress response, social dysfunction, and anhedonia, all core symptoms of depression, although to a greater extent in males than females at baseline. We have further shown that the “rewiring” of the brain that occurs due to ELA is caused by impaired microglial function, as indicated by diminished microglial process dynamics and synaptic pruning in males, but not females^4^. However, to our knowledge, no one has yet explored how ELA impacts peripheral immune markers with a focus on later risk for autoimmune diseases, which impact females more than males.

Here, our data demonstrate that <u>ELA during the first week of life</u> in mice significantly upregulates the level of IL-17A and additional autoimmunity-associated proteins in the <u>adult</u> blood, primarily in females. There were several proteins downregulated by ELA in both males and females, but these were primarily growth factors, rather than inflammatory cytokines or chemokines, which were upregulated specifically in females. Notably, autoimmune diseases like SLE have a higher prevalence in women than men, and there are known sex differences in immune function, with greater overall immune reactivity in women during adulthood^15^ .

Being a hallmark proinflammatory cytokine, IL-17A is found to increase in patients with SLE, an autoimmune disease that affects females and males at a ∼9:1 ratio, as well as in animals with lupus-like diseases in virtually all reports^10–12^. Furthermore, IL-17A is closely associated with other autoimmune diseases such as Sjögren’s syndrome, multiple sclerosis and rheumatoid arthritis, that also feature increased prevalence in females^16^ .

Analyzing each sex separately provided additional ELA-induced targets such as IL-13 and CCL3. Serum levels of IL-13 have been shown to associate with disease activity index of SLE (SLEDAI) and correlate with nerve growth factor (NGF) in lupus^17^. Recent data suggests CCL3^+^ monocytes are a commonly amplified immune subset in multiple autoimmune diseases including SLE and Sjögren’s syndrome^18^. Together, these data support the association between ELA-induced cytokine changes and autoimmune pathogenesis, particularly in females.

Limitations of the current work include a single measurement of proteomic changes in the blood of adult mice following rearing in ELA or control conditions. However, these data are derived from a highly sensitive NULISA assay to identify ELA-altered blood proteins in an unbiased manner, which highlighted IL-17A in particular as a potential mediator of ELA’s effects on autoimmune disease risk. Together, these data support our hypothesis that ELA alters inflammatory markers in adulthood that increase the risk for autoimmune diseases like lupus, and lay the foundation for future work to define the molecular mechanisms by which developmental programming of “immune memory” occurs after ELA and establish the impact of ELA on autoimmune pathogenesis.

## CRediT authorship contribution statement

**Madison M. Garvin**: Formal analysis, Investigation, Methodology, Project administration, Visualization, Writing-Review and Editing. **Ae R. Kim**: Formal analysis, Investigation, Methodology, Writing-Review and Editing. **Jessica L. Bolton:** Conceptualization, Formal analysis, Funding acquisition, Project administration, Supervision, Visualization, Writing-Original draft, Writing-Review and Editing. **Yun Liang:** Conceptualization, Formal analysis, Funding acquisition, Project administration, Supervision, Visualization, Writing-Original draft, Writing-Review and Editing.

## Acknowledgments

The authors would like to thank the Division of Animal Resources at Georgia State University for providing exceptional care to our animals.

## Data availability

Data will be made available on request.

